# Optimisation and validation of hydrogel-based brain tissue clearing shows uniform expansion across anatomical regions and spatial scales

**DOI:** 10.1101/423848

**Authors:** Adam L. Tyson, Ayesha Akhtar, Laura C. Andreae

## Abstract

Imaging of fixed tissue is routine in experimental neuroscience, but is limited by the depth of tissue that can be imaged using conventional methods. Optical clearing of brain tissue using hydrogel-based methods (e.g. CLARITY) allows imaging of large volumes of tissue and is rapidly becoming commonplace in the field. However, these methods suffer from a lack of standardised protocols and validation of the effect they have upon tissue morphology. We present a simple and reliable protocol for tissue clearing along with a quantitative assessment of the effect of tissue clearing upon morphology. Tissue clearing caused tissue swelling (compared to conventional methods), but this swelling was shown to be similar across spatial scales and the variation was within limits acceptable to the field. The results of many studies rely upon an assumption of uniformity in tissue swelling, and by demonstrating this quantitatively, research using these methods can be interpreted more reliably.

## Introduction

Fluorescence microscopy of fixed tissue sections is widely used in neuroscience, and biomedical science generally. However, light absorption (due to pigmentation) and scatter (due to heterogeneous refractive index (RI) of the tissue) limit the depth of tissue that can be imaged. To overcome this, tissue is usually sliced into thin sections (100 µm or less) which is laborious, and can introduce artefacts if large volumes of tissue are studied.

Light scatter due to lipid content is the predominant mechanism preventing deep imaging in brain tissue, and so tissue-processing methods have been developed to homogenise the RI of the tissue and reduce scatter. These methods are collectively known as tissue clearing, and were originally proposed a century ago^1^. More recently, the idea of tissue clearing for large-volume microscopy has been revisited. These methods have used different approaches, such as immersion in RI matching solutions^2^–^6^, the use of organic solvents^7^–^10^ and the direct removal of tissue lipids^11^–^13^. Of these, the methods relying on lipid removal, and particularly hydrogel-based methods (e.g. CLARITY^11^) have been those most adopted by the research community.

Hydrogel-based tissue clearing methods have so far been popular due to their reliability and flexibility (such as compatibility with antibody staining). Many variations on these methods have been published^11, 14–20^ but they all share a general core concept. Firstly, the tissue is incubated in a fixative solution containing paraformaldehyde (PFA) and acrylamide (with or without bis-acrylamide). This fixative binds biomolecules containing an amine group (chiefly proteins and nucleic acids) but not membrane phospholipids, and is then polymerised to to form a transparent hydrogel ‘matrix’ within the tissue. As the majority of lipids are not bound to this matrix, they can then be removed by using a detergent solution of sodium dodecyl sulfate (SDS) along with a combination of heat and electrophoresis or mechanical agitation to accelerate the process. Once the sample’s RI is matched using a high RI, low viscosity solution, the final result is a transparent and macromolecule permeable sample in which most protein and nucleic acid is preserved^11, 14, 20–22^.

There have been tremendous advances in tissue clearing along with imaging and analysis of large volumes of brain tissue. However, because these methods are not as mature as traditional methods (e.g. thin-section immunohistochemistry), two issues remain. The first is choosing an experimental protocol — there are many parameters to choose to ensure effective tissue clearing and staining. The second, and most important, is validation — these methods are starting to become routine, and yet there is very little information about how these methods affect tissue morphology.

Here we present an optimisation of a hydrogel-based tissue clearing and antibody staining protocol in adult mouse brain tissue. This was chosen as it is the most common, and most flexible use of tissue clearing in neuroscience. In addition, a detailed analysis was performed, comparing tissue morphology in cleared tissue to tissue processed using a more conventional method.

## Results

### Tissue clearing

To fully optimise hydrogel-based clearing of brain tissue, a number of parameters from the original report^11^ were varied. Samples were incubated whole, in hemispheres, or in slices taken using a brain slicing matrix^23^ and at room temperature or 37°C with or without shaking in clearing buffer (4% or 8% SDS) to clear. Clearing buffers were changed weekly, until the sample appeared visibly clear (i.e. until no distortion was visible when looking through the tissue (figure 1b).

**Figure 1.**
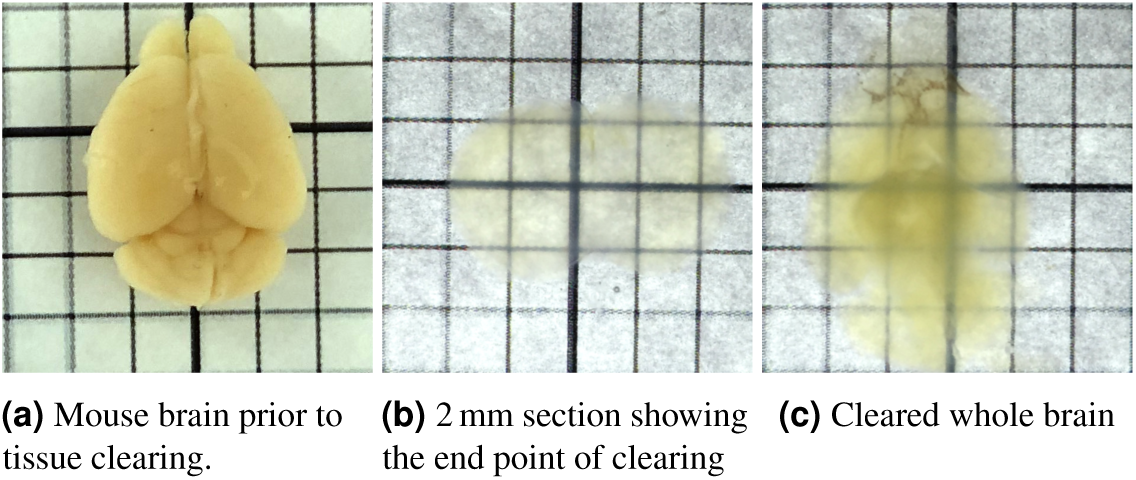
Mouse brain tissue incubated in hydrogel (A4B5P4), cleared using SDS and RI matched using 85 % glycerol.

All tissue was cleared successfully, showing little light scatter despite some discolouration (Fig 1b). Large volumes of brain tissue (e.g. whole, adult mouse brains, Fig 1c) could be cleared, despite more discolouration and scatter. There were no obvious differences in tissue clarity dependent upon hydrogel composition or the SDS concentration of the clearing buffer. An increase in SDS concentration (from 4% to 8 %) slightly increased the speed of tissue clearing, and shaking at 37 °C was required for complete, uniform clearance. The original (densest) hydrogel composition (A4B5P4) allowed 2 mm slices to clear in 5 weeks and whole adult mouse brains to clear in approximately 15 weeks. However, samples cleared much faster when prepared with lower-density hydrogel (Table 1). All hydrogel compositions allowed for successful staining, and so subjectively, hydrogel composition does not appear to affect antigen preservation. However, tissue rigidity is affected, and care must be taken not to damage samples prepared with a low-density hydrogel.

**Table 1.** Time taken to clear mouse brain hemispheres prepared with different hydrogel concentrations.

| Hydrogel composition | Density | Acrylamide [%] | Bis-acrylamide [%] | Paraformaldehyde [%] | Time taken for clearance [weeks] |
| --- | --- | --- | --- | --- | --- |
| A4B0P0 | Lowest | 4 | 0.00 | 0 | 3 |
| A1B5P1 | ↓ | 1 | 0.05 | 1 | 5 |
| A2B5P2 | ↓ | 2 | 0.05 | 2 | 6 |
| A4B5P4 | Highest | 4 | 0.05 | 4 | 12 |

### Tissue staining

#### Antibody staining

Following tissue clearing, a number of different antibodies (Fig 2, Supplementary Table 1) were tested, related to a variety of aspects of brain structure. Markers included cortical layer markers, striatal cell markers, inhibitory interneurons, synaptic markers, white matter markers and general cell type markers. Of these antibodies, nine produced reliable staining. These included the neuronal cell type markers CTIP2, CUX1, calbindin and parvalbumin along with the stains for neuronal projections (MBP and neurofilament) and general neuronal (NeuN) and astrocyte (GFAP) markers.

**Figure 2.**
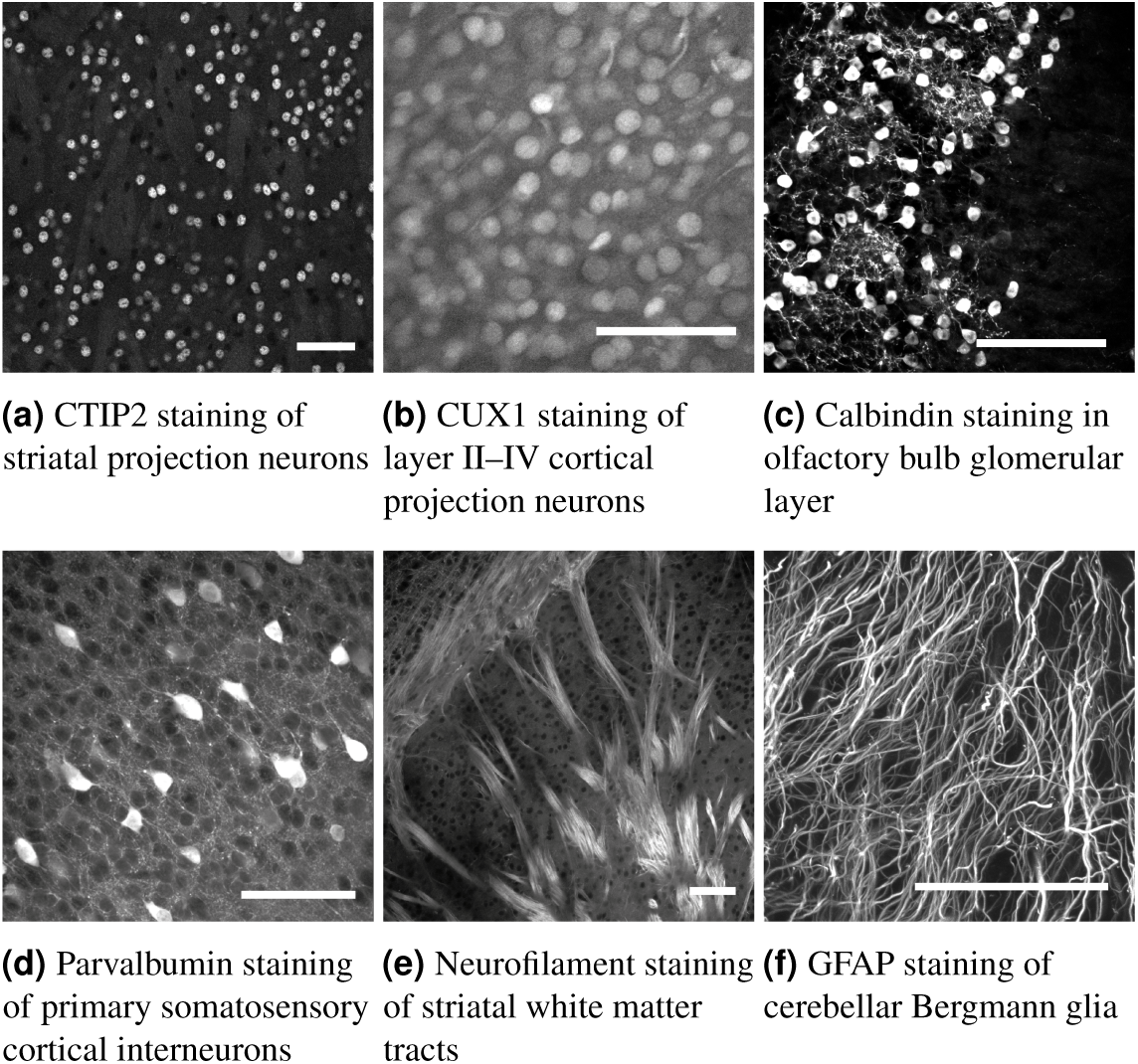
Examples of successful antibody stains in mouse brain tissue. Scale bars 100 µm.

#### Antibody penetration

To assess how much time antibodies took to diffuse into cleared tissue, the diffusion of the calbindin (rabbit) antibody was tested in mouse cortex (hydrogel composition — A4B5P4). This antibody was chosen as the protein is expressed relatively evenly across the cortex without being too dense (unlike neurofilament for example). Dense protein expression would complicate analysis, as antibody depletion would become the main factor limiting staining depth, rather than diffusion speed. To determine the speed of staining, the staining depth (the distance into the tissue at which brightly positive cells could be seen) was measured at four points in different areas of cortex. This was repeated for tissue samples stained for different lengths of time, and an average taken. The average antibody penetration is plotted in Fig 3.

**Figure 3.**
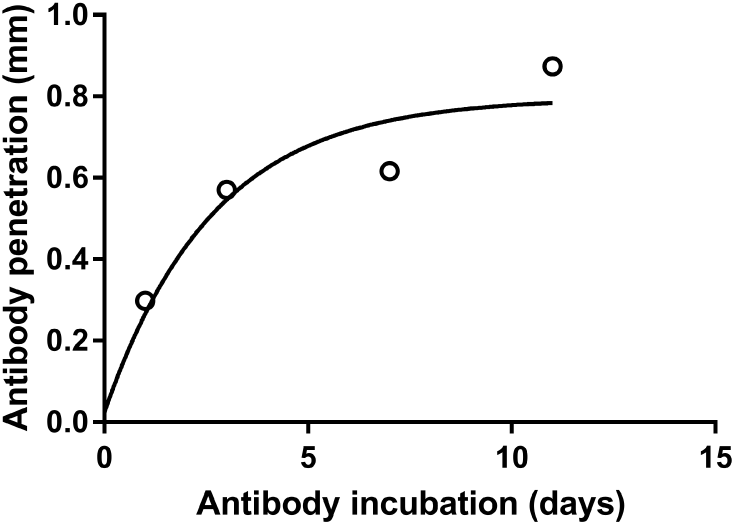
Antibody penetration depth as a function of incubation time, fitted with a single exponential.

#### Small molecule stains

Due to the slow diffusion of antibodies into cleared brain tissue, low molecular weight, non-antibody based fluorescent stains were investigated. These stains could all be used to successfully stain an entire, intact mouse brain within 24 hours, and as such greatly increase the flexibility of hydrogel-based tissue clearing. These dyes included nucleic acid, Nissl and myelin stains (Fig 4, Supplementary Table 2).

**Figure 4.**
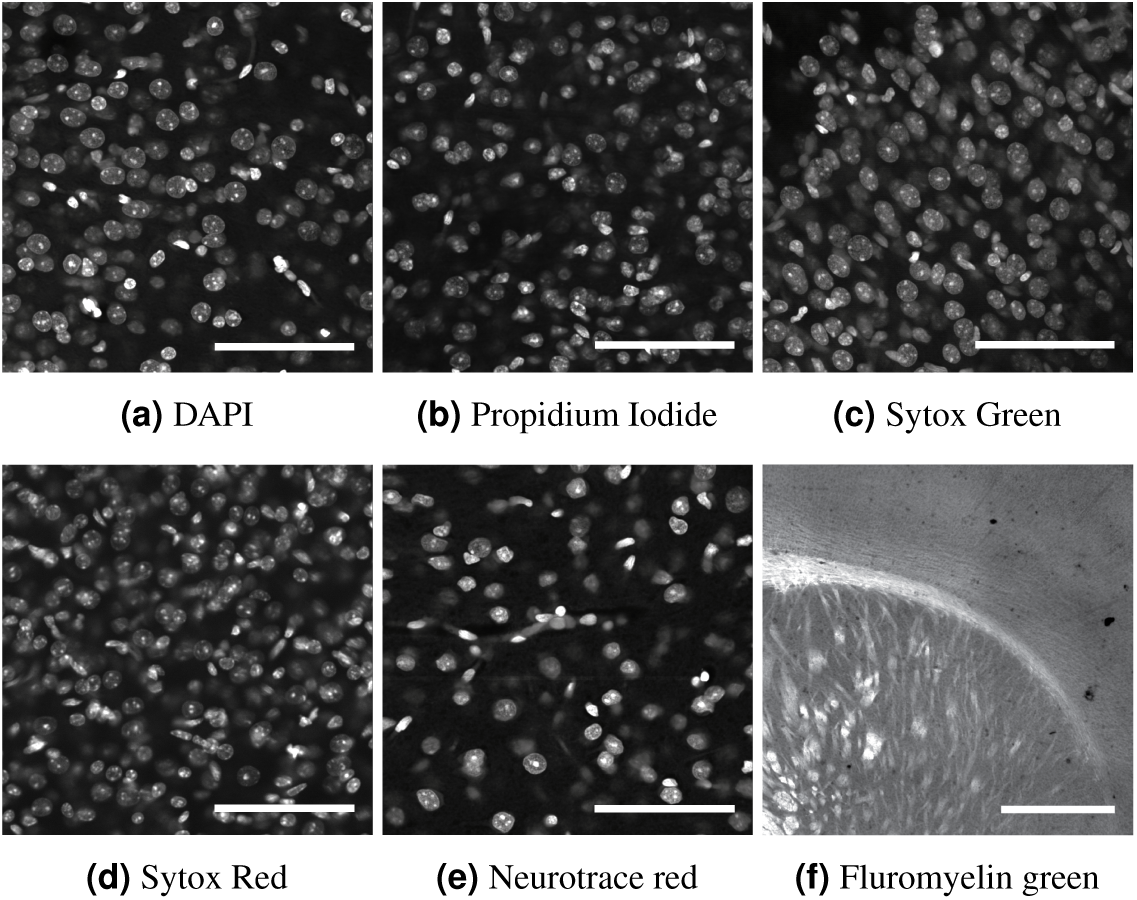
Other fluorescent stains in mouse brain tissue. All in primary somatosensory cortex and scale bars 100 µm other than fluromyelin (cortex and striatum scale bar 1 mm).

### Comparison to conventional fixation

To investigate the effects of tissue clearing, cleared tissue was compared to that fixed only using PFA and not cleared. Two adult, female littermate mice were perfused with phosphate buffered saline (PBS) and 4% PFA. One brain was post-fixed in hydrogel, before both brains were cut into 500 µm coronal sections and the hydrogel-fixed tissue was cleared. Following clearing, the cleared tissue and uncleared were treated identically and stained for a selection of markers. These were DAPI (nucleic acid), CTIP2 (cell body) and parvalbumin (whole cell). Firstly, two-dimensional (2D) images were acquired in a number of cortical and striatal areas to compare the cell density of CTIP2-and parvalbumin-positive cells. Two cleared and two uncleared slices were imaged for each cell type in each brain area, with 80 images taken of cortical parvalbuminergic cells (40 cleared and 40 uncleared), and 40 images taken of each cortical CTIP2 positive cells, striatal CTIP2-and striatal parvalbumin-positive cells. In all cases, the cell density was lower in cleared than uncleared tissue (*p*<0.001, Fig 5), suggesting tissue expansion. Although it is possible that the same results could occur owing to reduced antibody staining efficiency, this was thought to be unlikely, as the staining intensity of the positive cells was comparable between groups.

**Figure 5.**
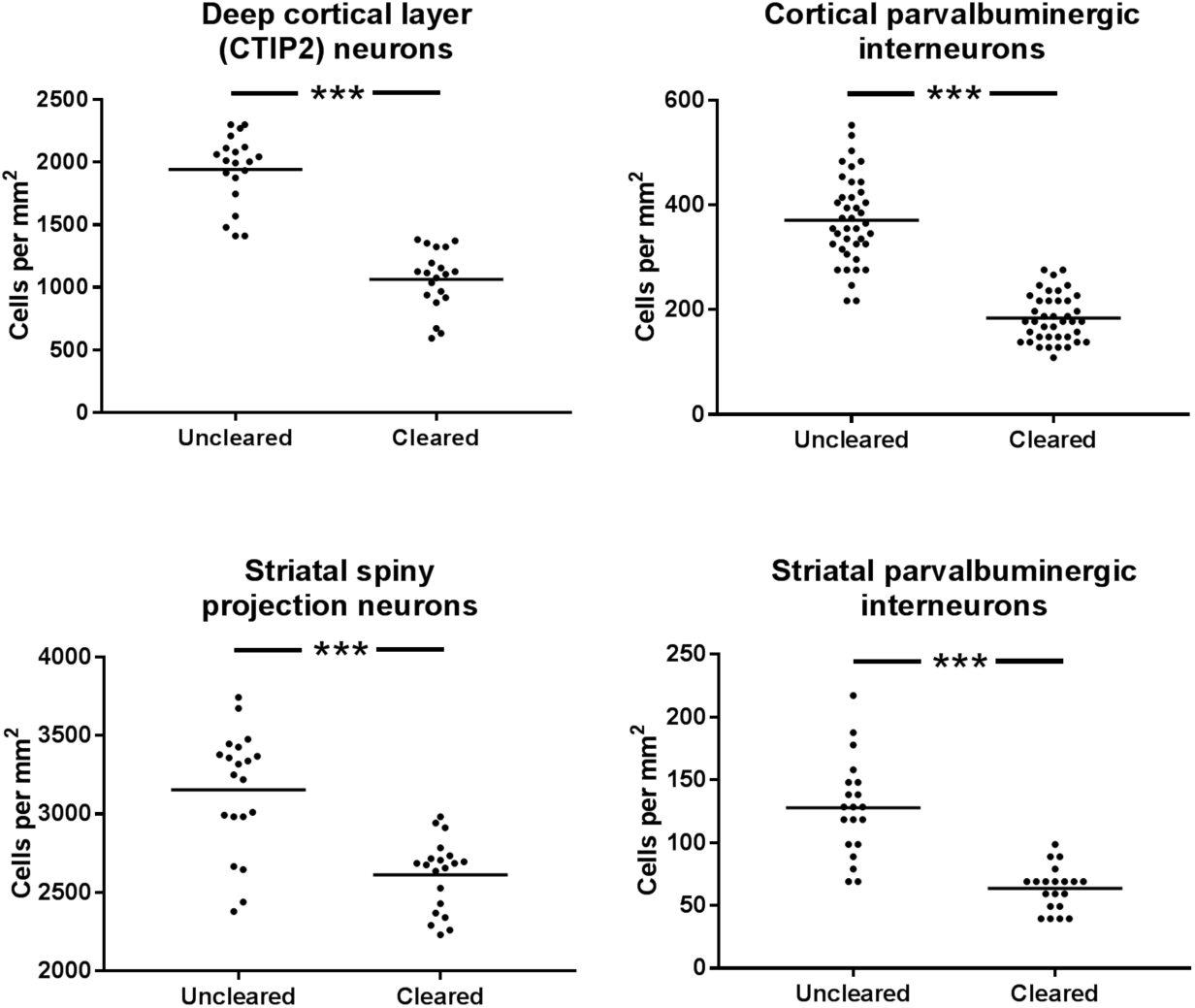
Cell counts of CTIP2 and parvalbumin positive cells in cortex and striatum, comparison between uncleared and cleared mouse brain tissue. Mean shown as a horizontal line.

To better understand whether these results were simply due to tissue expansion, and to assess whether this expansion occurred at different spatial scales, the volume of CTIP2-positive nuclei and parvalbumin-positive cells was investigated. The cell density result could have occurred due to uniform tissue expansion, or just expansion of the extracellular space. As before, two cleared sections and two uncleared sections were imaged for each cell type and each brain area. 30 cells in cleared tissue and 30 in uncleared tissue were imaged in three-dimensions (3D) for each cell type in each brain area, and the volumes (following manual segmentation) were compared between cleared and uncleared cells. In all cases, the cell volume was higher in cleared than in uncleared tissue (*p*<0.001, Fig 6).

**Figure 6.**
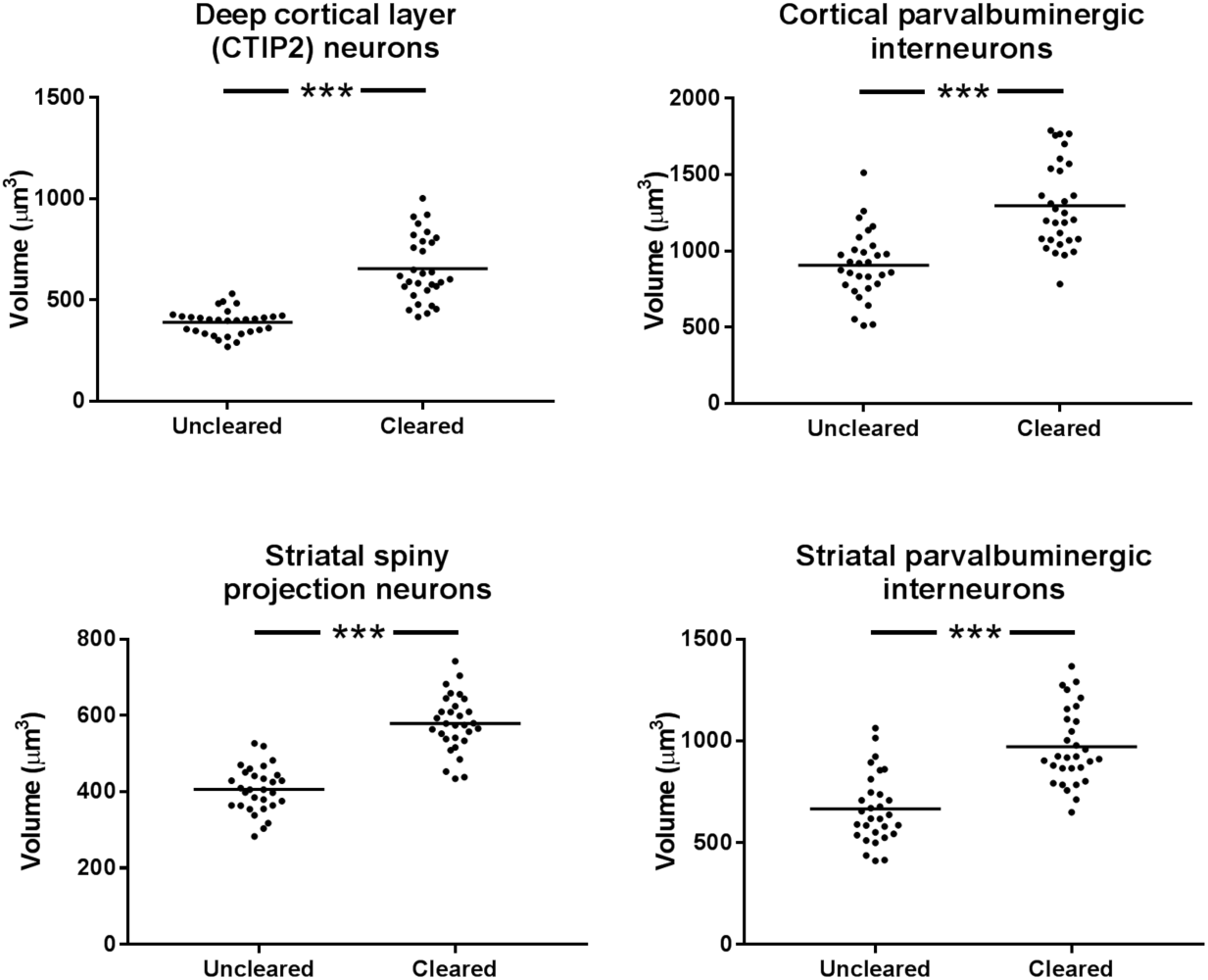
Volumes of CTIP2 and parvalbumin positive cells in cortex and striatum, comparison between uncleared and cleared mouse brain tissue. Mean shown as a horizontal line.

An increase in cell volume alongside a decrease in cell density suggested that the findings were due to general tissue expansion that is uniform at different spatial scales. As the tissue was already sectioned, it was impossible to look at the expansion of the entire brain, and so DAPI staining was used to measure cortical thickness. Cortical thickness was measured in two areas (motor cortex and barrel cortex) in nine brain hemispheres in each group (cleared and uncleared). The cortices of cleared tissue were thicker in both motor cortex (*p*=0.003) and barrel cortex (*p*=0.001, Fig 7), providing further evidence for general tissue expansion as the cause for the increased cell volume and reduced cell density.

**Figure 7.**
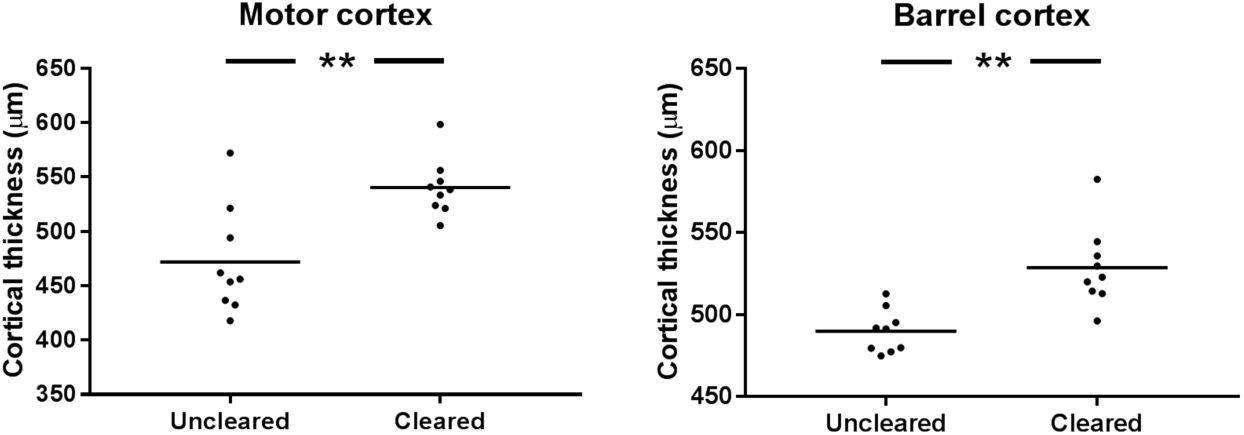
Comparison of cortical thickness in cleared and uncleared tissue, in motor and barrel cortices. Mean shown as a horizontal line.

Figures 5, 6 and 7 appear to show that tissue expansion affects all the measures in a similar fashion. These techniques are used under the assumption that any effects they have upon tissue structure are uniform across spatial scales and that no brain area or cell type is differentially affected. To assess whether this was the case, the variation in the effect of clearing upon the different parameters discussed above was calculated. Firstly, the relative change in the mean of each parameter was calculated, to give a measure of the expansion (Table 2). For volume or thickness measurements this was calculated as the cleared value divided by the uncleared value, and the inverse for the density measurements. This measure was then normalised to expansion in one dimension, so the 2D density measures were square-rooted, and the 3D volume measurements were cube-rooted. The coefficient of variance (CV, ratio of the standard deviation to the mean) of these measures was then calculated to give a metric (clearing CV) of how variable the effect of tissue expansion was upon the different measures of interest.

**Table 2.** Relative and adjusted expansion ratios forratios each individual measure comparing uncleared and cleared tissue.

|  | Measurement | Relative expansion metric | Adjusted ratio |
| --- | --- | --- | --- |
| Cell density | Cortical CTIP2 | 1.83 | 1.35 |
|  | Cortical Parvalbumin | 2.02 | 1.42 |
|  | Striatal CTIP2 | 1.21 | 1.10 |
|  | Striatal Parvalbumin | 2.00 | 1.41 |
| Cell volume | Cortical CTIP2 | 1.68 | 1.19 |
|  | Cortical Parvalbumin | 1.43 | 1.13 |
|  | Striatal CTIP2 | 1.42 | 1.12 |
|  | Striatal Parvalbumin | 1.46 | 1.13 |
| Cortical thickness | Motor cortex | 1.14 | 1.14 |
|  | Barrel cortex | 1.08 | 1.08 |

The nature of CVs is that there is no “rule of thumb” about what variance is high, low, acceptable or unacceptable. To evaluate the variance of the effect of clearing, the clearing CV was compared to acceptable levels of variance — the CVs of each of the individual measures in uncleared tissue. The variance of these measures in uncleared brain tissue is not considered to be a hindrance to detecting biological effects, and so if the clearing CV is of a similar magnitude, then tissue clearing can be thought of as acting uniformly in different brain areas, and at different spatial scales. The CV for tissue expansion was 0.110, which was lower than the CV for all the measures in uncleared tissue other than cortical thickness (0.104 & 0.027), as shown in Fig 8. Although this approach does not prove that clearing affects each metric uniformly, it suggests that the clearing CV is similar to generally accepted variance in the field.

**Figure 8.**
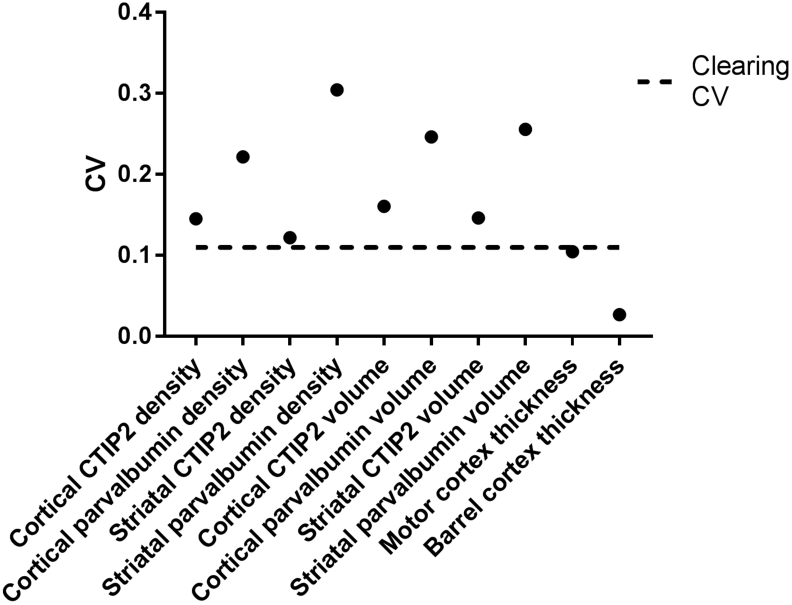
CVs for each measure in uncleared tissue with the CV due to clearing overlaid.

## Discussion

Hydrogel-based tissue clearing is very reliable method, but one issue is compatibility with antibody staining. We found that out of 22 antibodies tested, only nine were successful. However, these nine were very reliable and are likely to be of great use in the neuroscience community. The clearing process is a very harsh one which may cause antigen damage and prevent antibody staining, but this does not explain why some antibodies worked and not others. There is no obvious common feature of the successful antibodies, and this remains something to be studied further.

Another issue with antibody staining is the time taken for uniform staining with high signal-to-noise (SNR). Conventional primary and secondary antibody technology is not suitable for diffusion through large volumes of tissue. There are many ways to potentially speed this up, but possibly at the expense of SNR or tissue damage. These methods include using fluorophore-conjugated primary antibodies, low molecular weight single-domain antibodies^24^ or the use of electric fields to accelerate diffusion^25, 26^. Currently, none of these solutions are commercially available for most antibodies, and so researchers must rely on long incubation times of antibodies which practically limits the thickness of tissue that can be imaged.

The success of every small-molecule dye tested is potentially very useful. Cell nuclei stains have been used in conjunction with tissue clearing, but the Nissl and myelin stains have not yet (to our knowledge) been applied outside of method development. These stains can be used to stain much larger volumes of tissue than antibodies, and are much cheaper and easier to use. They are less specific than antibody stains, but at large tissue volumes they represent a great increase in specificity and resolution compared to competing techniques (e.g. magnetic resonance imaging).

We show that tissue clearing using a hydrogel-based approach appears to swell the brain tissue in a uniform way. The variability of tissue expansion upon different measures in different brain areas is generally less than the natural variability associated with these measures in uncleared tissue. For this reason, it is not thought that tissue clearing will introduce any biases that could affect the interpretation of any results. It is also well known that many tissue fixatives (including PFA) can shrink brain tissue considerably^27^, so it may be that the swelling introduced by clearing compared to PFA-only fixation may return the tissue closer to the structure found *in vivo*.

Since the original description of hydrogel-based tissue clearing^11^, there have been close to a hundred publications applying and extending the method^28^, the majority of them in neuroscience. We have described a simple and reliable protocol allowing investigation of many common neuroanatomical features including cell densities, white matter and glial structure along with the morphology of a number of specific cell types. Very few of the existing papers have measured the effect upon morphology of these techniques, other than just describing tissue expansion very broadly. We have shown that although these methods do cause tissue expansion (at least compared to existing techniques), it can be thought of as happening uniformly across spatial scales. Therefore as long as these methods are applied consistently, the results from them can be interpreted as reliably as with other, more established methods.

## Methods

### Tissue clearing

All procedures were performed under local King’s College London Animal Welfare and Ethical Review Body approval and under UK Home Office project and personal licenses, where necessary, in accordance with the Animals (Scientific Procedures) Act 1986. C57BL/6J mice were euthanised by cervical dislocation, and brain tissue was rapidly dissected and incubated in ice cold hydrogel solution. Hydrogel solutions were made up in PBS with 0.25% VA-044 photoinitiator (Wako Chemicals GmbH, DE) according to Table 1. Samples were incubated (with shaking) at 4 °C for one week. To prevent inhibition of the polymerisation by oxygen, the samples were degassed using nitrogen^11^ then polymerised in a 37 °C water bath for 3 hours. Excess hydrogel was removed, was incubated in clearing solution (SDS in in 0.2M boric acid, pH=8.5) whole, as hemispheres, or in 500 µm (using a vibratome, VT1000S, Leica Biosystems GmbH, Germany) or 2 mm (using a brain-slicing matrix^23^) sections. Clearing buffer was exchanged after 24 and 48 hours, and then weekly until samples became transparent. Samples were then washed in PBSTN_3_ (0.1 % Tx100 and 0.01 % sodium azide in PBS) and stored until staining.

### Tissue staining

For antibody staining, samples were incubated with primary antibody made up in PBSTN_3_ with gentle shaking at 37 °C for 7 days. Secondary antibody staining was performed as per the primary with the appropriate AlexaFluor 488 conjugated secondary antibody (Life Technologies Ltd, UK) at a concentration of 1:50. Small molecule stains were also carried out in PBSTN_3_, but at room temperature (approximately 20 °C), and for 24 hours. All samples were thoroughly washed in PBSTN_3_ and then incubated in 85% glycerol in PBS for 24 hours prior to imaging.

### Imaging and data analysis

Antibody and dye optimisation imaging was carried out on a variety of confocal and multiphoton microscopes. The comparison between cleared and uncleared tissue was carried out on a Nikon Eclipse 80i C1 laser-scanning confocal microscope (Nikon Instruments Europe BV, NL) using the parameters in Supplementary Table 3. All images in this section were 512 × 512 pixels and taken with a 150 µm pinhole.

All images were analysed using ImageJ^29–32^. Cell densities were determined manually and cortical thickness measurements were taken by measuring the length of straight lines drawn across the cortex. To measure cell volumes, cells were segmented manually using the segmentation editor plugin^33^ and the volumes of the ROIs were calculated using the 3D ImageJ suite^34^. All image figures were generated using ImageJ and all statistics and plots were generated using Prism (Graphpad Software Inc, USA). In all cases, datapoints represent regions of interest, and not individual animals. Distributions were assessed for normality using the Shapiro-Wilk test and all were normal other than cortical CTIP2 cell counts in uncleared tissue. For ease of interpretation, all comparisons were carried out using the parametric t-test. Due to the notable general reduction in variance for some variables in cleared tissue, Welch’s adaptation was used throughout (**= *p* <0.01,***= *p* <0.001). Full test statistics are shown in Supplementary Table 4.

## Acknowledgements

This work was supported by a Guy’s and St. Thomas’ Charity Prize PhD scholarship to ALT. We acknowledge financial support from the Innovative Medicines Initiative Joint Undertaking under grant agreement no. 115300, resources of which are composed of financial contribution from the European Union’s Seventh Framework Programme (FP7/2007–2013) and EFPIA companies’ in kind contribution, the Mortimer D Sackler Foundation and the Sackler Institute for Translational Neurodevelopment (ALT and LCA).

## Author contributions statement

ALT and LCA conceived the experiments, ALT and AA conducted the experiments and analysed the results, ALT and LCA wrote the manuscript. All authors reviewed the manuscript.

## Additional information

### Data availability

All raw data generated in the production of this manuscript is available on request.

## SUPPLEMENTARY INFORMATION

**Table 1.**
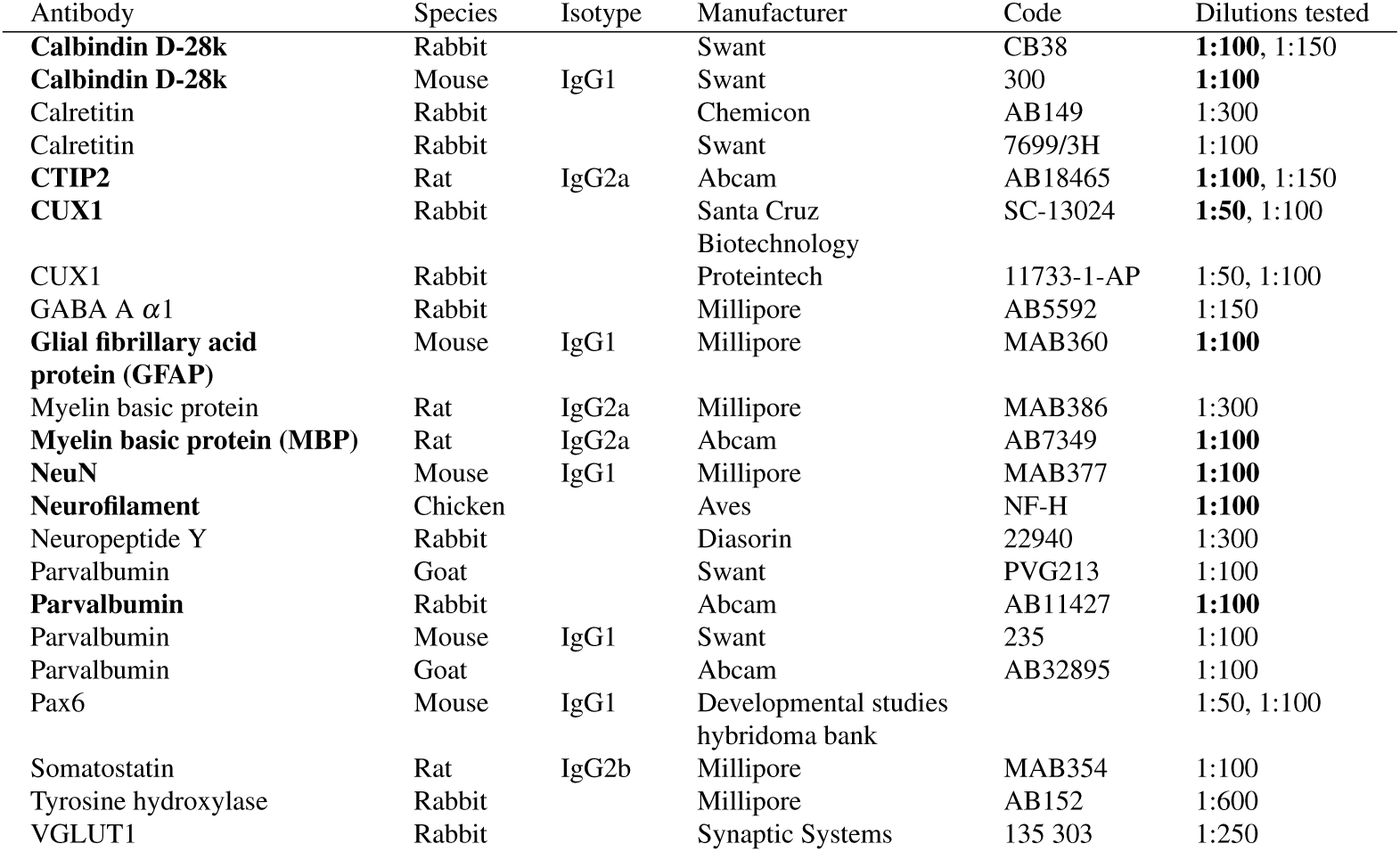
Primary antibodies tested, successful antibodies and dilutions in bold.

**Table 2.** Fluorescent small molecule dyes tested.

| Stain | Target | Successful concentrations |
| --- | --- | --- |
| DAPI | Nucleic acids (cell nuclei) | 3.6 μM to 36 μM |
| Propidium iodide | Nucleic acids (cell nuclei) | 3 μM |
| Sytox green | Nucleic acids (cell nuclei) | 5 μM |
| Sytox red | Nucleic acids (cell nuclei) | 5 μM |
| Neurotrace red | Nissl bodies (neurons) | 1:100 |
| Fluoromyelin green | Myelin (white matter) | 1:100 |

**Table 3.**
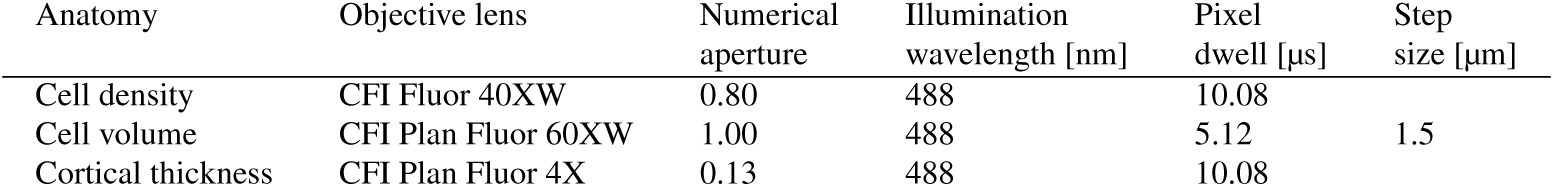
Tissue clearing comparison microscopy acquisition parameters.

**Table 4.**
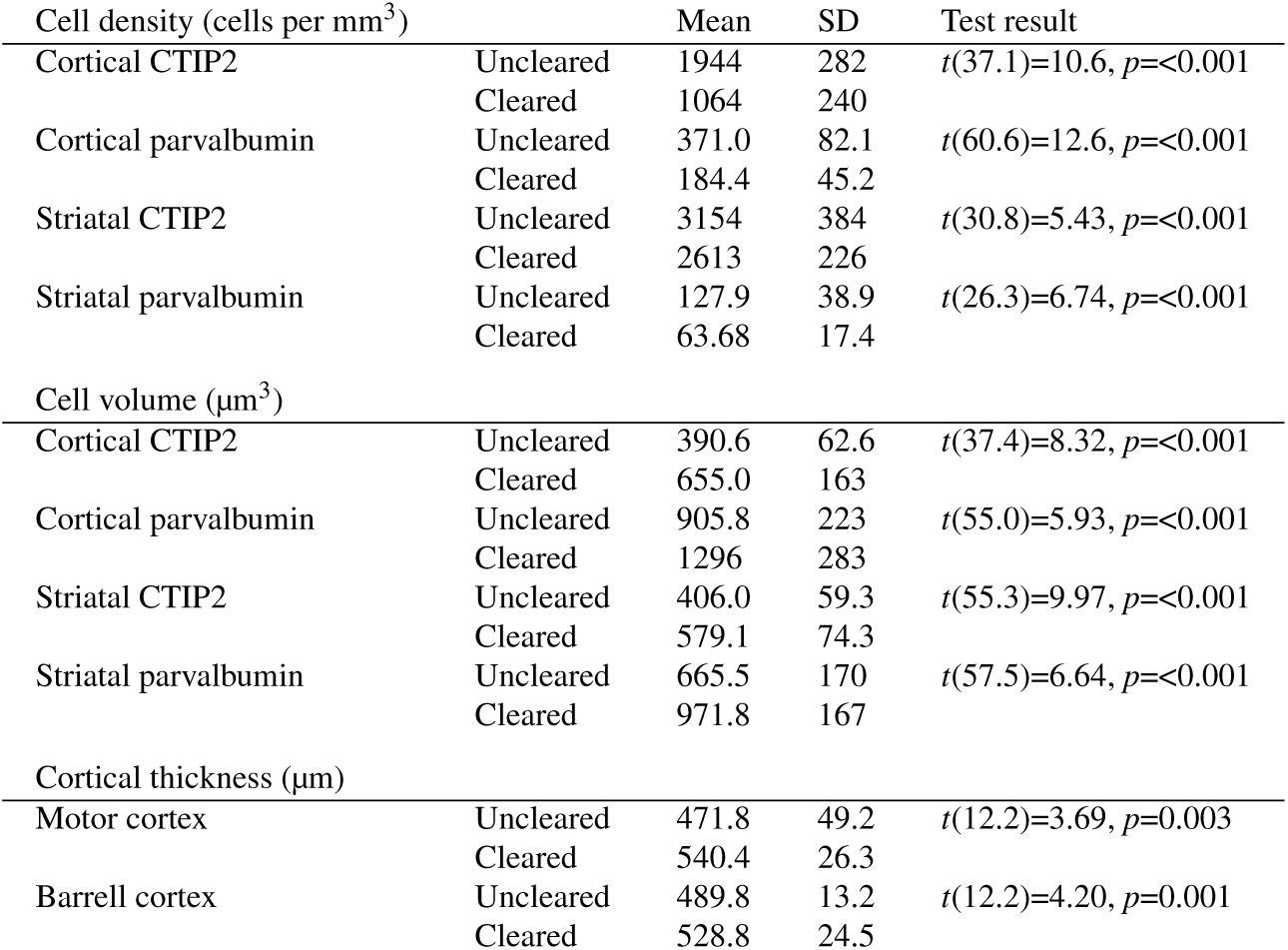
Descriptive statistics and *t*-test results of the comparison between uncleared and cleared tissue.

