## Supplemental tables for "Optimisation and validation of hydrogel-based brain tissue clearing shows uniform expansion across anatomical regions and spatial scales"

### SUPPLEMENTARY INFORMATION

| Antibody | Species | Isotype | Manufacturer | Code | Dilutions tested |
|---|---|---|---|---|---|
| **Calbindin D-28k** | Rabbit | | Swant | CB38 | **1:100**, 1:150 |
| **Calbindin D-28k** | Mouse | IgG1 | Swant | 300 | **1:100** |
| Calretitin | Rabbit | | Chemicon | AB149 | 1:300 |
| Calretitin | Rabbit | | Swant | 7699/3H | 1:100 |
| **CTIP2** | Rat | IgG2a | Abcam | AB18465 | **1:100**, 1:150 |
| **CUX1** | Rabbit | | Santa Cruz Biotechnology | SC-13024 | **1:50**, 1:100 |
| CUX1 | Rabbit | | Proteintech | 11733-1-AP | 1:50, 1:100 |
| GABA A $\alpha$1 | Rabbit | | Millipore | AB5592 | 1:150 |
| **Glial fibrillary acid protein (GFAP)** | Mouse | IgG1 | Millipore | MAB360 | **1:100** |
| Myelin basic protein | Rat | IgG2a | Millipore | MAB386 | 1:300 |
| **Myelin basic protein (MBP)** | Rat | IgG2a | Abcam | AB7349 | **1:100** |
| **NeuN** | Mouse | IgG1 | Millipore | MAB377 | **1:100** |
| **Neurofilament** | Chicken | | Aves | NF-H | **1:100** |
| Neuropeptide Y | Rabbit | | Diasorin | 22940 | 1:300 |
| Parvalbumin | Goat | | Swant | PVG213 | 1:100 |
| **Parvalbumin** | Rabbit | | Abcam | AB11427 | **1:100** |
| Parvalbumin | Mouse | IgG1 | Swant | 235 | 1:100 |
| Parvalbumin | Goat | | Abcam | AB32895 | 1:100 |
| Pax6 | Mouse | IgG1 | Developmental studies hybridoma bank | | 1:50, 1:100 |
| Somatostatin | Rat | IgG2b | Millipore | MAB354 | 1:100 |
| Tyrosine hydroxylase | Rabbit | | Millipore | AB152 | 1:600 |
| VGLUT1 | Rabbit | | Synaptic Systems | 135 303 | 1:250 |

**Table 1.** Primary antibodies tested, successful antibodies and dilutions in bold.

| Stain | Target | Successful concentrations |
|---|---|---|
| DAPI | Nucleic acids (cell nuclei) | 3.6 µM to 36 µM |
| Propidum iodide | Nucleic acids (cell nuclei) | 3 µM |
| Sytox green | Nucleic acids (cell nuclei) | 5 µM |
| Sytox red | Nucleic acids (cell nuclei) | 5 µM |
| Neurotrace red | Nissl bodies (neurons) | 1:100 |
| Fluromyelin green | Myelin (white matter) | 1:100 |

**Table 2.** Fluorescent small molecule dyes tested.

| Anatomy | Objective lens | Numerical aperture | Illumination wavelength [nm] | Pixel dwell [µs] | Step size [µm] |
|---|---|---|---|---|---|
| Cell density | CFI Fluor 40XW | 0.80 | 488 | 10.08 | |
| Cell volume | CFI Plan Fluor 60XW | 1.00 | 488 | 5.12 | 1.5 |
| Cortical thickness | CFI Plan Fluor 4X | 0.13 | 488 | 10.08 | |

**Table 3.** Tissue clearing comparison microscopy acquisition parameters.

| Cell density (cells per $mm^3$) | | Mean | SD | Test result |
|---|---|---|---|---|
| Cortical CTIP2 | Uncleared | 1944 | 282 | $t(37.1)=10.6$, $p=<0.001$ |
| | Cleared | 1064 | 240 | |
| Cortical parvalbumin | Uncleared | 371.0 | 82.1 | $t(60.6)=12.6$, $p=<0.001$ |
| | Cleared | 184.4 | 45.2 | |
| Striatal CTIP2 | Uncleared | 3154 | 384 | $t(30.8)=5.43$, $p=<0.001$ |
| | Cleared | 2613 | 226 | |
| Striatal parvalbumin | Uncleared | 127.9 | 38.9 | $t(26.3)=6.74$, $p=<0.001$ |
| | Cleared | 63.68 | 17.4 | |
| Cell volume ($\mu m^3$) | | | | |
| Cortical CTIP2 | Uncleared | 390.6 | 62.6 | $t(37.4)=8.32$, $p=<0.001$ |
| | Cleared | 655.0 | 163 | |
| Cortical parvalbumin | Uncleared | 905.8 | 223 | $t(55.0)=5.93$, $p=<0.001$ |
| | Cleared | 1296 | 283 | |
| Striatal CTIP2 | Uncleared | 406.0 | 59.3 | $t(55.3)=9.97$, $p=<0.001$ |
| | Cleared | 579.1 | 74.3 | |
| Striatal parvalbumin | Uncleared | 665.5 | 170 | $t(57.5)=6.64$, $p=<0.001$ |
| | Cleared | 971.8 | 167 | |
| Cortical thickness (µm) | | | | |
| Motor cortex | Uncleared | 471.8 | 49.2 | $t(12.2)=3.69$, $p=0.003$ |
| | Cleared | 540.4 | 26.3 | |
| Barrell cortex | Uncleared | 489.8 | 13.2 | $t(12.2)=4.20$, $p=0.001$ |
| | Cleared | 528.8 | 24.5 | |

**Table 4.** Descriptive statistics and *t*-test results of the comparison between uncleared and cleared tissue.
